# Restoration of Direct Corticospinal Communication Across Sites of Spinal Injury

**DOI:** 10.1101/546374

**Authors:** Naveen Jayaprakash, David Nowak, Erik Eastwood, Nicholas Krueger, Zimei Wang, Murray G. Blackmore

**Affiliations:** Department of Biomedical Sciences, Marquette University, 53201

## Abstract

Injury to the spinal cord often disrupts long-distance axon tracts that link the brain and spinal cord, causing permanent disability. Axon regeneration is then prevented by a combination of inhibitory signals that emerge at the injury site and by a low capacity for regeneration within injured neurons. The corticospinal tract (CST) is essential for fine motor control but has proven refractory to many attempted pro-regenerative treatments. Although strategies are emerging to create relay or detour circuits that re-route cortical motor commands through spared circuits, these have only partially met the challenge of restoring motor control. Here, using a murine model of spinal injury, we elevated the intrinsic regenerative ability of CST neurons by supplying a pro-regenerative transcription factor, KLF6, while simultaneously supplying injured CST axons with a growth-permissive graft of neural progenitor cells (NPCs) transplanted into a site of spinal injury. The combined treatment produced robust CST regeneration directly through the grafts and into distal spinal cord. Moreover, selective optogenetic stimulation of regenerated CST axons and single-unit electrophysiology revealed extensive synaptic integration by CST axons with spinal neurons beyond the injury site. Finally, when KLF6 was delivered to injured neurons with a highly effective retrograde vector, combined KLF6/NPC treatment yielded significant improvements in forelimb function. These findings highlight the utility of retrograde gene therapy as a strategy to treat CNS injury and establish conditions that restore functional CST communication across a site of spinal injury.

**Significance Statement:** Damage to the spinal cord results in incurable paralysis because axons that carry descending motor commands are unable to regenerate. Here we deployed a two-pronged strategy in a rodent model of spinal injury to promote regeneration by the corticospinal tract, a critical mediator of fine motor control. Delivering pro-regenerative KLF6 to injured neurons while simultaneously transplanting neural progenitor cells to injury sites resulted in robust regeneration directly through sites of spinal injury, accompanied by extensive synapse formation with spinal neurons. In addition, when KLF6 was delivered with improved retrograde gene therapy vectors, the combined treatment significantly improved forelimb function in injured animals. This work represents important progress toward restoring regeneration and motor function after spinal injury.

## Introduction

Spinal cord injury disrupts the exchange of information between the brain and distal cord, causing impairments in sensory, motor, and autonomic function. In cases of incomplete injury, severed axons often sprout spontaneously to form new connections with spared tracts, creating detour circuits that re-route information around the injury (1–6). Recent work involving transplants of neural progenitor cells has also succeeded in creating novel relay circuits as host axons invade and innervate graft-derived neurons, which in turn extend lengthy axons that innervate neurons in the caudal spinal cord (7–9). These indirect circuits, both endogenous and graft-derived, have yielded some gains in motor function after injury(4, 6, 7, 9, 10). Recovery remains partial, however, even as various pharmacological, rehabilitation, and stimulation strategies have attempted to enhance their functional output (4, 6). Fundamentally, in the face of supraspinal control systems that evolved to rely on direct connectivity between supraspinal nuclei and spinal neurons, there may be a limit to the ability of detour or relay circuits to replace lost function, particularly for tasks involving fine motor control. Thus, to complement progress in creating indirect replacements circuitry after injury, there remains a pressing need to restore the ability of supraspinal neurons to communicate directly with distal spinal neurons.

Doing so likely requires regenerative growth by injured axons. Axon regeneration in the central nervous system (CNS) is blocked by dual barriers: a low neuron-intrinsic capacity for axon growth in adults (11, 12) and an inhospitable environment around the injury site (13–15). Corticospinal tract (CST) neurons, important mediators of fine motor control in both rodents and humans, have historically regenerated poorly but more recently have responded, albeit partially, to several pro-regenerative treatments (16, 17). The neuron-intrinsic growth ability of CST neurons can be enhanced by deleting growth-inhibitory genes (18, 19) or forcing the expression of pro-regenerative factors (20, 21). A transcription factor called KLF6 has emerged as a particularly potent promoter of CST axon growth by activating a set of complementary gene networks involved in axon extension (22). Yet these neuron-intrinsic interventions only partially restore growth ability, and stimulated axons generally circumvent partial injuries, but fail to traverse complete injuries, suggesting persistent environmental inhibition (23). To address extrinsic inhibition, an important advance has been the demonstration that grafts of neural progenitor cells (NPCs) placed into sites of spinal injury can attract regenerative ingrowth from CST axons (10). Nevertheless, one major limitation is that a majority of regenerating CST axons extend only about 1mm into the grafts, and rarely re-enter distal host tissue caudal to the injury site (10). This finding hints that even when extrinsic inhibition is neutralized with improved tissue, the low growth ability of CST axons limits full growth.

Here we test a novel combinatorial strategy to enhance CST regeneration by simultaneously addressing intrinsic and extrinsic barriers to growth. Using a murine model of deep cervical injury, we combine KLF6 gene delivery and NPC grafting to evoke robust CST regeneration that extends completely through injuries and into distal host tissue. Both forelimb and hindlimb populations of CST axons participated in the regenerative growth. Optogenetic stimulation of regenerated axons and single unit recordings of spinal neurons confirmed the ability of CST axons to evoke spinal activity distal to injury sites, indicating restoration of direct communication between cortex and spinal tissue distal to the injury. Finally, by harnessing a newly developed retrograde AAV vector to improve gene delivery, we found that combined KLF6 expression and NPC grafting significantly improved fine motor control in a horizontal ladder task and pellet retrieval task. These data indicate that combined NPC grafting and delivery of pro-regenerative transcription factors can restore direct neural communication across sites of spinal injury.

## Results

### Combined KLF6 expression and NPC grafts enhance CST regeneration

KLF6 is a pro-regenerative transcription factor that is expressed by corticospinal tract (CST) neurons during periods of axon growth, and then downregulated with maturation (22, 24). We showed previously that forced re-expression of KLF6 in adult CST neurons enhances compensatory CST sprouting and regenerative growth after partial spinal injuries (22). Importantly, however, this KLF6-stimulated growth occurred through spared grey matter. Here we tested whether KLF6 over-expression can stimulate CST regeneration through a deeper and more challenging injury. AAV9-KLF6 or AAV9-EBFP control, mixed with AAV-EYFP tracer, were injected to the left motor cortex of adult C57Bl/6J mice. Animals then received an injury to cervical spinal cord in which a wife-knife was inserted at the midline, the blade extended toward the right edge of the spinal cord and lowered/raised in three successive cycles to a depth of 1.1mm. In contrast to the prior injuries, this produced a more tear-like injury, affecting the majority of grey matter (**Supp. Fig. 1**). Eight weeks later, sections of spinal cord were prepared, and CST growth was quantified as the number of EYFP+ axons that intersected virtual lines at set distances below the injury, normalized to total EYFP+ axons counted in the medullary pyramids. As expected, AAV-EBFP-treated animals showed minimal CST growth below the injury site (**Supp. Fig. 2A,B**). In contrast to prior findings (22), KLF6-treated animals displayed only a non-significant trend toward enhanced CST growth in this injury paradigm (**Supp. Fig. 2C,D)**. These data indicate that when confronted with a deeper and more severe injury, KLF6-stimulated axons are largely unable to regenerate, a finding reminiscent of prior findings with closely related KLF7 (21, 23). Thus, although KLF-based manipulations elevate intrinsic regenerative ability in CST axons (22), extrinsic barriers continue to constrain growth.

**Fig. 1.**
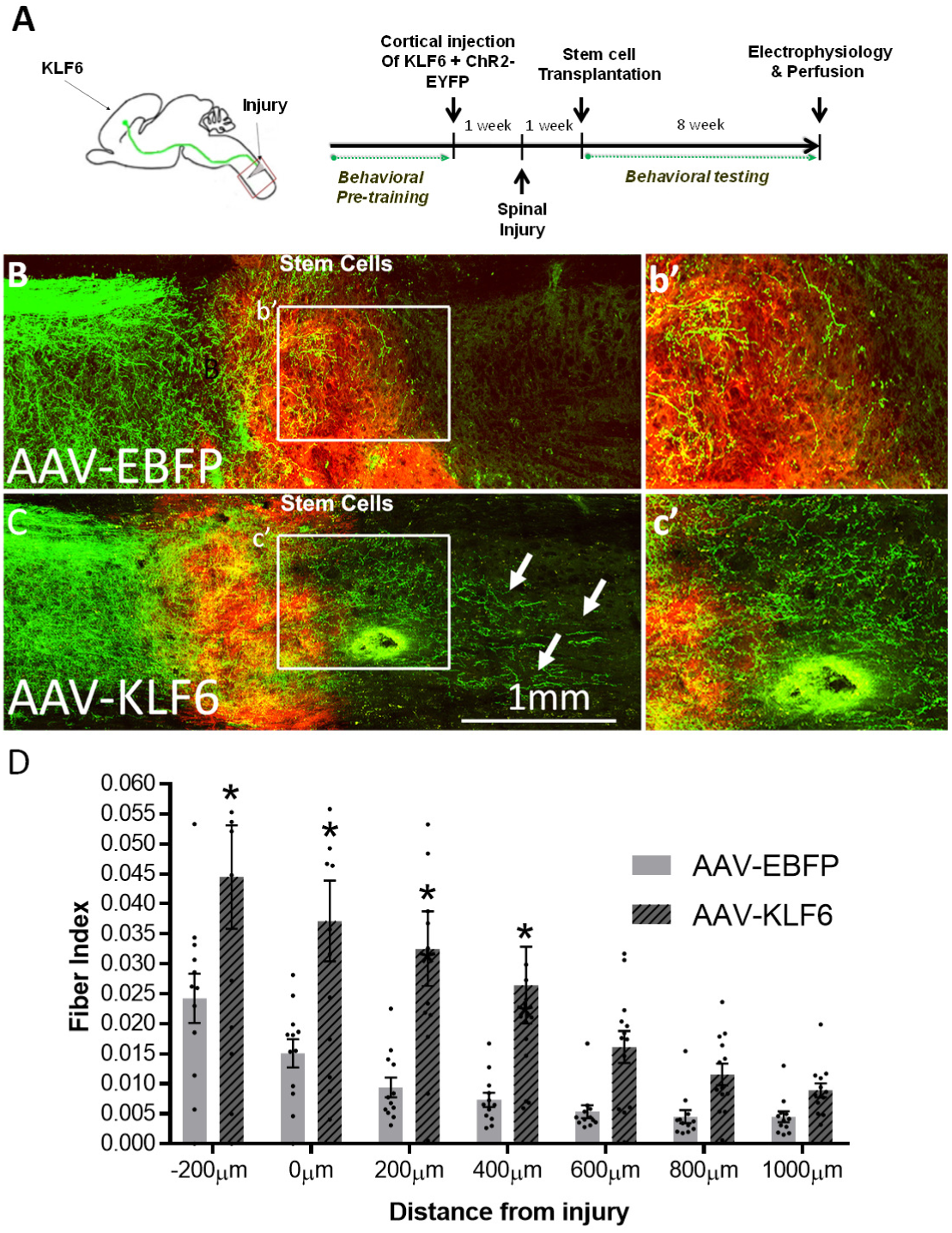
Combined KLF6 expression and NPC grafting improves CST regeneration. (A) Adult mice received cortical injection of AAV-ChR2-EYFP and AAV-KLF6 or AAV-EBFP control, followed by cervical hemisection and transplantation of NPCs labeled with tdTomato. (B-E) Eight weeks after NPC grafting, CST axons (EYFP+, green) regenerate into NPC grafts (red). In animals treated with KLF6, CST axons extend beyond the distal graft/host border and into spinal tissue caudal to the lesion (arrows, C). (D) Quantification of the number of axonal profiles that extend beyond the injury shows a significant elevation of CST regeneration in KLF6-treated animals. N=12 per group, *p<.05, repeated measures ANOVA.

**Fig. 2.**
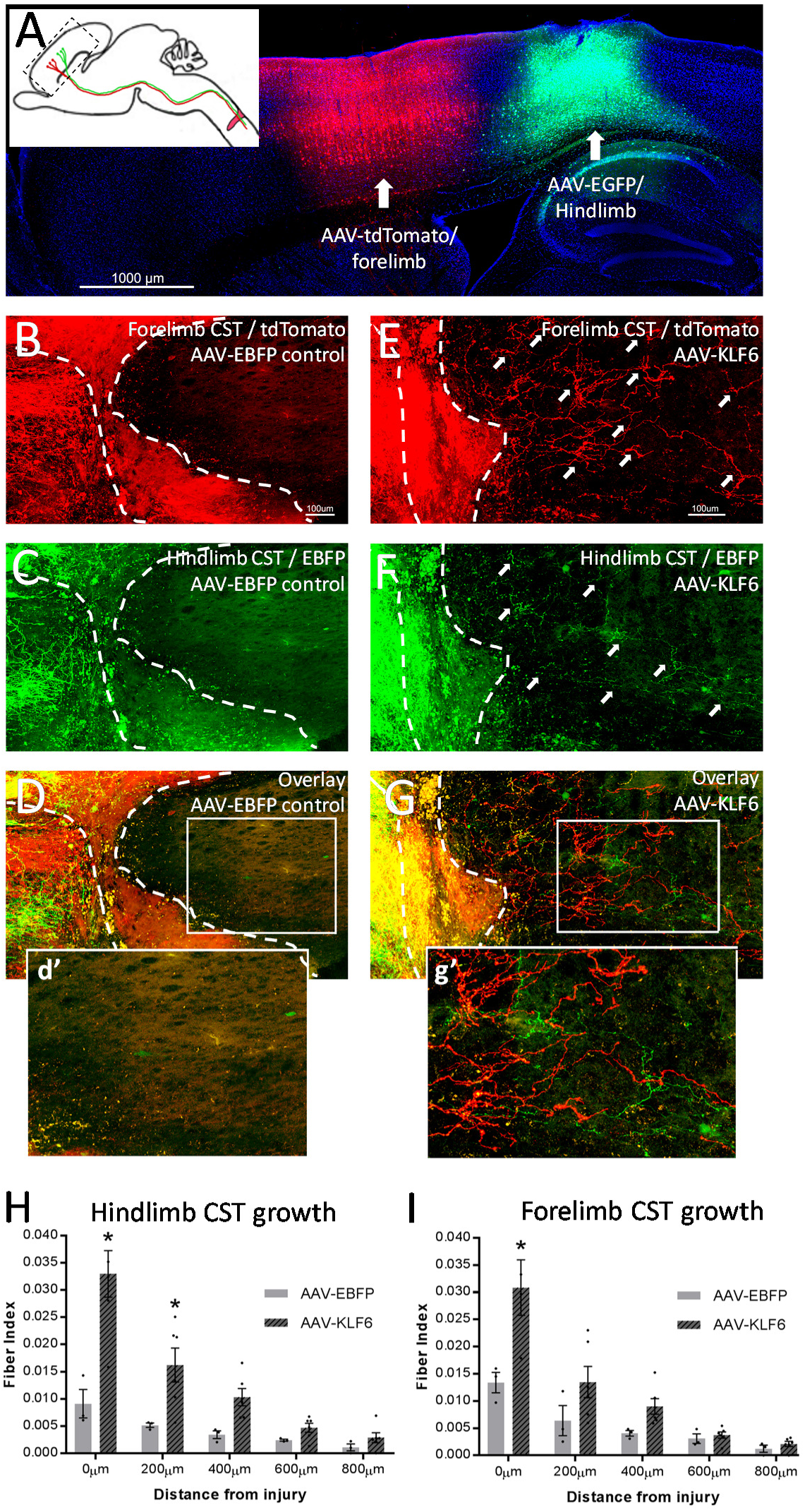
Selective targeting of forelimb or hindlimb CST populations (A) was achieved by focally injecting into respective motor cortex regions. Axon regeneration from hindlimb CST with EBFP control vector (B, H) showed little growth (<0.010 fiber index) caudal to injury site. KLF6 over-expression in hindlimb CST significantly enhanced growth phenotype at injury and 200µm caudal to injury site (p<0.05, two-way ANOVA) (E, H). Forelimb CST axons with EBFP control vector showed little (<0.015 fiber index) growth beyond injury site, while KLF6 treated animals had significantly more axon growth at the injury site only (*p<0.05, repeated measures ANOVA) (F, I). With control vector (EBFP) both forelimb (EGFP) and hindlimb (tdTomato) axon populations (D) show little growth past injury (d’). KLF6 overexpression in both forelimb (td-Tomato) and hindlimb (EGFP) axon populations (G) significantly improves regeneration past the injury (g’). N= 6 KLF6, 3 CTR, axon regeneration from forelimb and hindlimb CST was not significantly different (*p>0.05, Two way ANOVA, Tukey’s multiple comparison)

We therefore employed grafts of embryonic spinal cells supported by a fibrin matrix and growth factors, shown previously to create growth-permissive interfaces with host tissue and to support ingrowth by injured CST axons (7, 8, 10). To confirm the ability of these grafts to attract regenerative CST growth, adult mice received cortical injection of control AAV-EBFP and AAV-ChR2-EYFP tracer and deep wire knife injury, followed one week later by transplantation of cells from E12.5 embryonic spinal cord, labeled by transgenic tdTomato expression (**Fig. 1A**). Eight weeks after transplantation, examination of spinal sections showed consistent survival and integration of grafts and ingrowth by in CST axons (EYFP+), confirming the grafts’ suitability as a growth substrate (**Supp. Table 1**). Consistent with prior reports, however, most of this CST growth penetrated less than 1mm into the grafts, and rarely traversed the entire graft to re-enter distal host tissue (10). Thus, the availability of growth-permissive embryonic tissue restores axon regeneration by CST neurons only partially, likely reflecting continued neuron-intrinsic constraints to growth. To simultaneously improve both neuron-intrinsic and -extrinsic conditions for growth, adult mice received cortical injection of AAV-KLF6 and AAV-ChR2-EYFP tracer, followed by spinal injury and NPC grafting as described above. Eight weeks later, animals that received combined KLF6/NPCs showed a significant increase in CST growth into graft tissue beyond injury sites (**Fig. 1D**). In addition, the number of axons that extended past the distal graft boundary and re-entered host tissue was also significantly elevated (**Fig. 1D**). Importantly, numerous axons were detected traversing the grafted tissue and re-entering host spinal cord, indicating substantial growth directly through the original injury site, as opposed to circumventing the injury through spared tissue (**Fig. 1C)**. These data indicate that when injured axons are provided with growth-supportive tissue, KLF6 expression enhances regenerative growth by CST neurons.

### KLF6 promotes CST regeneration from both forelimb and hindlimb CST

Distinct sub-regions of cortex project axons to distinct spinal cord levels and control the movement of different joints (25) Notably, CST input to cervical and lumbar spinal cord arise from separate regions of cortex (e.g. “forelimb” and “hindlimb” cortex). To test KLF6’s effects across distinct cortical sub-regions, we focally injected AAV-KLF6 or AAV-EBFP control to forelimb or hindlimb cortex, and then performed injury and NPC grafting as previously. In half of the animals, forelimb CSTs were labeled by co-injection of AAV-EGFP and hindlimb by AAV-tdTomato, and in other half the color scheme was reversed. Eight weeks after transplantation, animals were sacrificed, and axon growth quantified in spinal sections, normalized to total transduced axons as above. Importantly, total counts of EGFP and tdTomato-traced axons were similar within treatments, ruling out systematic effects of tracer detection. In control animals, both forelimb and hindlimb CST axons extended into grafts, but rarely into distal tissue. In contrast, KLF6-treated axons traversed grafts and re-entered distal tissue, with similar amounts of growth arising from forelimb and hindlimb populations. These data indicate that KLF6’s ability to promote axon growth is robust across different sub-regions of cortex and consistent across experiments, highlighting its versatility.

### Regenerated CST axons form functional synapses on distal spinal neurons

To test the ability of regenerated CST axons to form effective synapses we paired selective optogenetic stimulation of CST terminals with single-unit recording from spinal neurons in intact animals. As described above, animals received mixed cortical injection of AAV-ChR2-EYFP and AAV-EBFP or AAV-KLF6, followed by cervical injury. Half of the animals then received NPC grafts. Eight weeks later animals were anesthetized and their spinal cords exposed between C3 and C6. 473nm light was focally directed to locations up to 1000µm above the injury site, to the center of the injury, and up to 1000µm distal to the injury. In this way optical stimulation specifically triggers synaptic release from CST terminals, which uniquely express ChR2 in spinal tissue. At each location, a 32-multichannel electrode was inserted 1000µm into the spinal cord, with sensitivity to activity from individual cell bodies but not fibers of passage (26). Spontaneously active units were identified and the average change in firing during light exposure was calculated for each unit. Units that displayed significant increases in firing during light stimulation (p<.001, paired t-test; see methods) were classified as CST-responsive. Overall, this strategy quantifies the ability of CST activity to alter firing in spinal neurons, and thus probes the potential for restored communication.

As expected, all treatment groups displayed post-synaptic responses when light stimulation was directed above the injury site, where intact CST axons were present (Fig 3A-D). These data serve as a positive control for the ability optogenetically stimulated CST axons to drive post-synaptic activity. In locations distal to the injury, animals that received no NPC grafts, regardless of KLF6 treatment, displayed minimal post-synaptic responses, consistent with the low level of CST ingrowth to these regions. Thus, as expected, spinal injury reliably interrupts communication between CST axons and caudal spinal neurons. Animals that received NPC grafts and EBFP gene treatment displayed moderate response to CST stimulation at the injury site (11% of units classified as CST-responsive). Importantly, because NPC grafts were centered in this location, the enhanced firing rate during optical stimulation likely reflected synaptic integration of CST axons with graft-derived neurons, consistent with prior findings (10). Distal to the injury, 10% and 7% of host neurons responded to CST stimulation at 500um and 1mm, respectively. These data indicate that host CST axons, which enter grafts and occasionally exit distally, likely form functional synapses on both graft-derived and host neurons, albeit with modest frequency. Finally, in animals that receive both grafts and KLF6 treatment, a substantially larger fraction of units responded to CST stimulation both within and beyond grafts. Indeed, 34% and 29% of spontaneously active spinal units responded to CST activation at positions 500um and 1000um beyond the spinal injury, respectively. In addition, in CST-responsive units the average change in firing rate below the injury was significantly larger in KLF6 than in EBFP-treated animals (KLF6: 16.9 (±2.1 SEM) spikes/s, EBFP: 6.0 (±0.6 SEM) spikes/s p=.0068, 2 tailed t-test). Thus KLF6 treatment of CST axons produces substantial elevation of direct neural communication distal to a site of spinal injury.

**Fig. 3.**
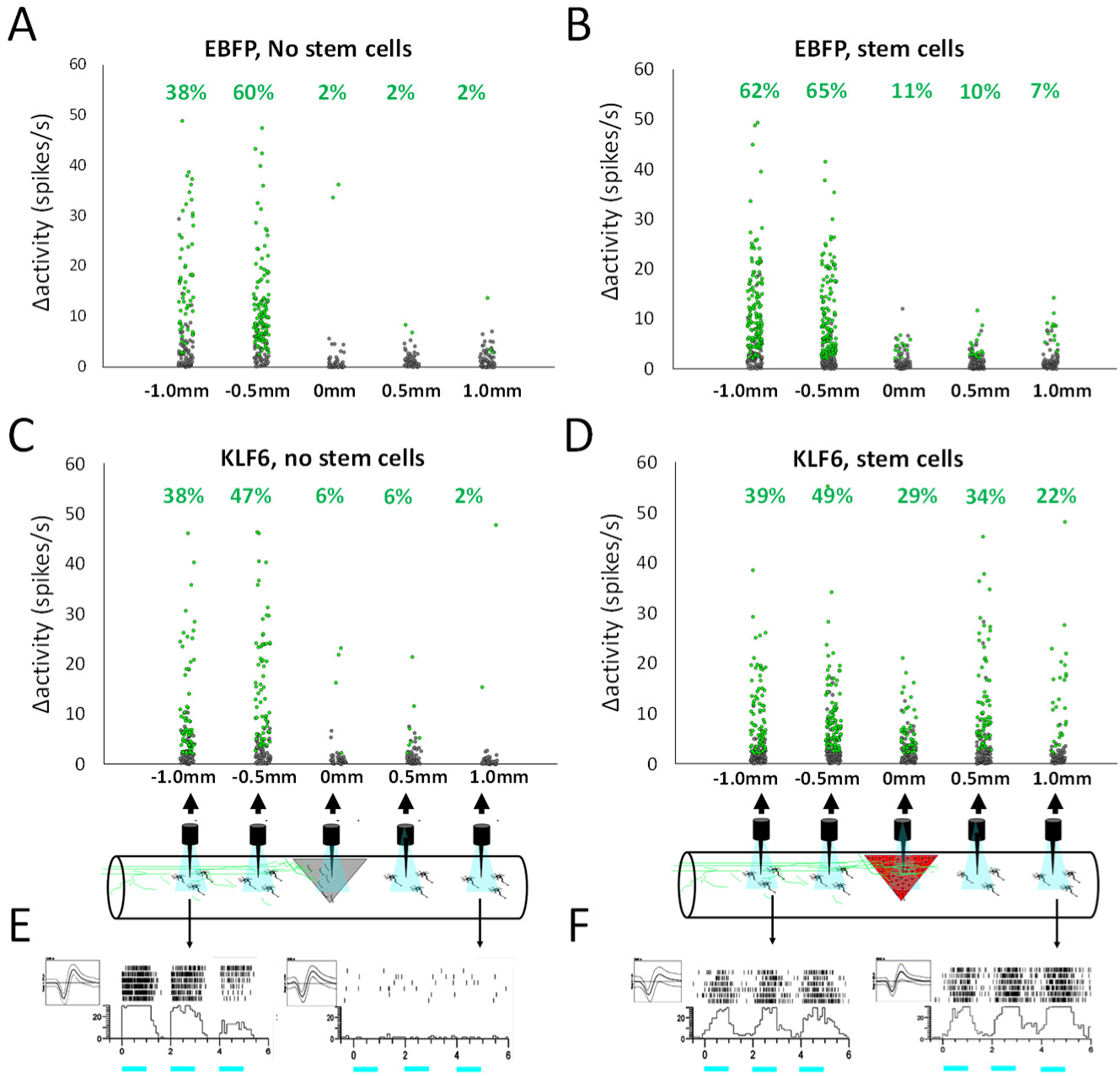
Selective optogenetic stimulation and single unit recording reveal synaptic connectivity between regenerated CST axons and spinal neurons. Adult mice received cortical injection of AAV-ChR2-EYFP and AAV-KLF6 or AAV-EBFP control, followed by cervical hemisection and NPC transplants or saline control. Eight weeks later, 473nm light was focally delivered to sites rostral to, within, and caudal to the injury site while single-unit recordings were performed with a 32-channel electrode. ChR2 is present only on CST axons, thus enabling CST-selective stimulation. (A-D) Each dot represents a single spinal unit, with position indicating the average change in firing rate during light stimulation and green indicating significance (p<.001, paired t-test). In the absence of NPCs, light-responsive cells were detected rostral to the injury, but <6% responded at any position caudal to the injury (A,C). (B) NPC grafts alone yielded up to 10% response rate caudal to the injury. (D) In animals that received both NPCs and AAV-KLF6, 34% and 22% of spinal units responded to CST stimulation at 500µm and 1000 µm caudal to the injury, respectively. (E,F) show representative raster histograms and waveforms of spinal units.

### Retrograde KLF6 delivery and NPC grafting promote forelimb function after cervical spinal injury

To test for potential restoration of forelimb function, animals treated with NPC grafts and/or KLF6 were tested on a horizontal ladder task at four and eight weeks post-injury. As expected, spinal injuries significantly increased the frequency of misplaced steps by the affected forelimb. NPC grafts produced a non-significant trend toward improved forelimb placement at both four and eight weeks, and KLF6 expression produced no significant change (**Supp. Fig. 3A,B**). Thus, despite the enhanced axon growth and synaptic connectivity from combined KLF6/NPC grafts, this treatment did not yield behavioral improvement in this task. We hypothesized that behavioral recovery could be limited in part by the fact that forelimb movement is controlled by CST neurons spread broadly across sensorimotor cortex, in coordination with subcortical populations including the rubrospinal and retriculospinal tracts (3, 27, 28). Thus focal delivery of KLF6 by injections to the cortex may not reach a broad enough population of supraspinal neurons. To deliver KLF6 more broadly we employed Retro-AAV, a newly developed vector with enhanced retrograde properties. We recently showed that delivery of Retro-AAV to the cervical spinal cord produces highly efficient transduction of CST neurons throughout the cortex, in addition to subcortical populations including the rubrospinal tract (29). To confirm the ability of Retro-AAV to drive KLF6 expression we co-injected Retro-AAV2-KLF6 and Retro-AAV2-tdTomato to cervical spinal cord. Two weeks later, upregulation of KLF6 was readily detectable in western blot analysis of whole cortex, and by immunohistochemistry in CST and rubrospinal tract neurons (**Fig. 4 A-D**). We therefore repeated behavioral assessment of animals treated with combined KLF6 and NPCs, this time delivering KLF6 by injection of Retro-AAV to the spinal cord at the time of NPC grafting. Cortical tracing in a subset of animals confirmed extensive regeneration by CST axons in the combined treatment group (**Fig. 4E-H**). Combined Retro-KLF6 and NPC grafting produced a significant reduction in mistargeted steps on the horizontal ladder task and a significant increase in the number of pellets that were displaced and retrieved by the right forelimb (**Fig. 4I, J**). Thus, in the presence of growth-permissive tissue grafts, KLF6 delivery with a retrograde vector produces significant improvements in forelimb function after spinal injury.

**Fig. 4.**
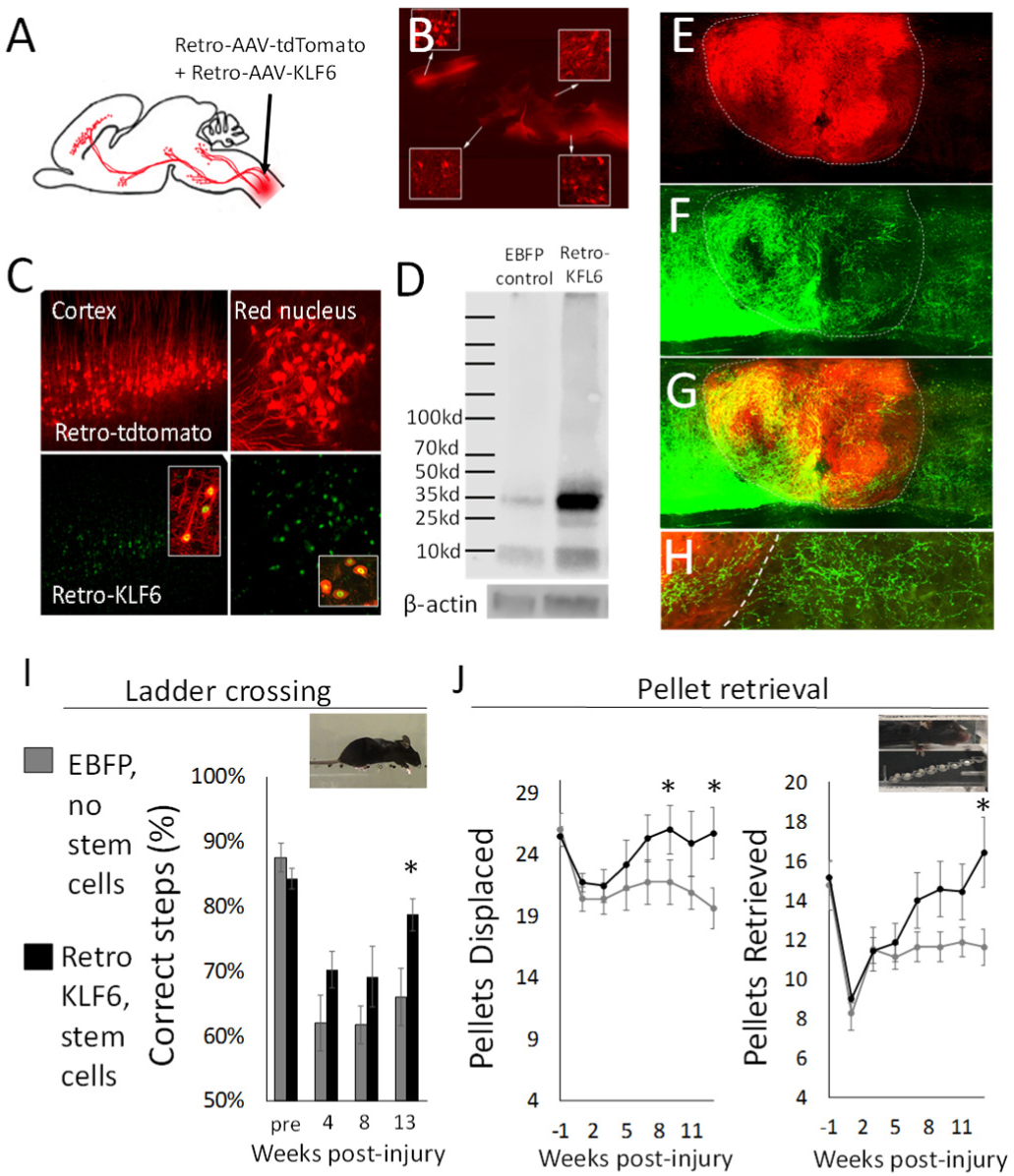
Widespread delivery of KLF6 with retrograde vectors, combined with NPC grafts, improves forelimb function after cervical spinal injury. (A-D) Adult mice received cervical injection of AAV2-Retro-KLF6 with AAV2-Retro-tdTomato tracer. Two weeks later, tdTomato fluorescence confirmed widespread retrograde transduction (B) and immunohistochemistry confirmed successful overexpression of KLF6 in retrogradely transduced cells (C). (D) Western blotting confirmed elevated KLF6 expression in cortical tissue two weeks after retrograde delivery to cervical spinal cord. (E-H) Adult mice received cervical hemisection injury and cervical injection of AAV2-Retro-KLF6, followed one week later by NPC transplants. Twelve weeks later, AAV9-EGFP was injected to cortex to trace CST axons. (E-H) Similar to anterograde KLF6, retrogradely expressed KLF6 produced robust CST growth into and beyond NPC grafts. (I,J) Compared to control animals that received EBFP and no NPCs, animals treated with AAV2-Retro-KLF6 and NPC grafts displayed significant improvements in forelimb placement on the

## Discussion

We have shown that following spinal injury, regeneration of the corticospinal tract is substantially enhanced by simultaneously expressing a pro-regenerative transcription factor in injured neurons and supplying growth-permissive tissue bridges to injured axons. Injured CST axons traversed the grafted tissue and re-entered distal host tissue, where they established synaptic connections. This outcome was consistent across multiple experiments and was observed in both forelimb and hindlimb-directed populations of CST neurons. Moreover, when delivered with a retrograde vector, KLF6 combined with NPC grafts produced significant restoration of forelimb function in cervically injured mice. These findings establish conditions that reliably evoke functional regeneration by corticospinal tract neurons across a site of spinal injury.

The promotion of CST growth after spinal injury has been a long-standing goal in regeneration research. In work spanning several decades, a great diversity of cell types have been transplanted into animal models of spinal injury in an effort to provide CST axons with a more growth-permissive tissue environment (30–38). Grafted cells have succeeded in eliciting growth from other supraspinal populations (e.g. propriospinal, raphespinal, bulbospinal, and sometimes rubrospinal), but CST axons often respond minimally or not at all (30–36). Thus, compared to other populations, CST neurons appear to possess an intrinsically low capacity for growth and/or more stringent requirements for extrinsic growth substrates. In this context, it is highly significant that NPC grafts, supported by growth factors and a fibrin/thrombin matrix, were recently shown to act as suitable substrates for CST axon growth (10). Here we confirm the ability of NPC grafts to attract growth from injured CST axons; the reproducibility of this effect across research labs marks important progress toward the long-sought goal of evoking regenerative growth from this critical population.

Also consistent with prior observations, however, we found CST growth through the grafts to be incomplete, mostly reaching only the proximal half of the graft and very rarely extending to distal host tissue (10). One important implication of this pattern of growth is that to provide functional improvements, NPC grafts alone are unable restore the original CST circuitry but instead must act by creating novel relay circuits through grafted neurons. There is now substantial evidence that new relay circuits do form in grafts, based on the growth and synaptogenesis by host neurons into grafts and exuberant extension of graft-derived axons into distal host tissue (7, 8). Moreover, recent findings indicate that these relay circuits contribute functionally to improvements in hindlimb stepping motions(9). There may be limits, however, to the ability of these relay circuits to restore motor control, particularly for fine forelimb movements normally controlled by the CST. Timing and synchrony of firing between neural circuits is essential for both the learning and execution of fine motor tasks (39). From a temporal perspective, relay circuits in NPC grafts create a built-in delay for movement and the return of sensory information, greatly complicating coincidence detection. Spatially, as host axons invade the graft they appear to terminate in a highly disorganized pattern that lacks clear topography (8, 10, 40), although there is some evidence that some populations do select appropriate cell types as synaptic targets (40). Again, this initial disorganization complicates any engagement of Hebbian processes to sculpt effective movement (41, 42). It is notable that despite the robust growth in and out of NPC grafts and the existence of new relay connections, improvements even in stereotyped motions such as locomotion remain only partial(7, 9).

To address this constraint, we harnessed forced expression of a pro-regenerative transcription factor to drive substantial regeneration of CST axons completely through NPC grafts and into distal host tissue. KLF factors comprise a 17-member family that play widespread roles in development, including both positive and negative regulation of axon growth (24, 43). KLF6 and the closely related KLF7 stand out as the only members known to be pro-regenerative (21, 22, 24). In contrast to KLF7, which required addition of a VP16 transcriptional activation domain to evoke CST regeneration (21), KLF6 effectively promotes CST axon growth in its wild type form(22). Here we showed that when environmental constraints are relieved by cell transplants into the site of injury, CST neurons with forced expression of KLF6 extended axons through sites of spinal injury and back into distal host tissue. CST regeneration was consistent across experiment and between cortical regions, highlighting the potential of combined intrinsic and extrinsic-targeted treatments to reliably evoke CST regeneration. Importantly, direct optical stimulation of these regenerated axons produced robust post-synaptic responses from host neurons in distal cord, indicating the re-establishment of direct synaptic connectivity. Thus, this combined approach reliably yielded a neural substrate for direct communication between the cortex and distal spinal cord, directly through a site of spinal injury.

Finally, a key finding was that delivery of KLF6 by retrograde vectors, but not direct cranial injection, improved forelimb function after spinal injury. We recently characterized retrograde transduction after spinal injections of AAV2-Retro, and showed widespread transduction of CST neurons throughout the cortex, in addition to subcortical nuclei that participate in skilled movements, including the red nucleus and reticular formation (29). The likely explanation for the difference in behavioral outcomes is that compared to focal delivery to subregions of motor cortex, retrograde vectors deliver therapeutic transgenes to a larger number and greater diversity of supraspinal cell types. Importantly, retrograde gene delivery also offers substantial advantages over direct cortical injection from a translational perspective. Retrograde injections could be readily combined with existing cell transplantation procedures, and in human patients a small number of injections to spinal targets could potentially reach widespread supraspinal populations in a way that is highly impractical using direct cranial injections. Thus, these data highlight the potential for retrograde KLF6 vectors to supplement current grafting approaches and further improve functional outcomes after spinal injury.

## Materials and Methods

### Injury and AAV delivery

Spinal cord injury was performed between cervical region C4-C5 using a wire knife to create a lesion of 1.1mm depth. AAV9-KLF6 and AAV-EBFP were created by PCR-based cloning into a pAAV-MCS vector, followed by production at the Univ. of N. Carolina vector core, as described previously (22). AAV9-CamKII-ChR2(H134R)-EYFP was obtained from the Univ. of North Carolina Viral Vector Core. pAAV-CAG-tdTomato (codon diversified) was a gift from Edward Boyden (Addgene viral prep # 59462-AAV5). Vectors were delivered to five locations spanning the motor cortex of adult female C57/Bl6 mice, (1×10^13^ particles/ml, 0.3μl each) using a Kopf stereotaxic instrument, Hamilton syringe, and Stoelting QSI infusion pump.

### NPC preparation and transplantation

Spinal cords from tdTomato-expressing (Jax 007676) E12 embryos were dissociated by trituration in 1X HBSS buffer and resuspended in Neurobasal medium. 1 million cells were injected by Picospritzer 5 to five locations surrounding the injury site in thrombin / fibrinogen and BDNF, VEFF, bFGF and MDL28170VEGF (BDNF 50ng/µl, bFGF 10ng/µl, VEGF 10ng/µl, MDL28170 50µM)

### Behavior

As described previously (26) mice were pre-trained on a horizontal ladder (30cm long, 1cm rung spacing) and staircase pellet apparatus (Lafayette Instruments, Lafayette IN) until performance plateaued, then tested every four weeks (ladder) or bi-weekly (retrieval) after injury.

### In vivo optogenetics and electrophysiology

Extracellular recording were obtained using a multichannel silicon electrode inserted 1000 µ m from the surface of the spinal cord. 473 nm laser was focused via collimator on the point of electrode insertion to stimulate terminal firing of ChR2-expressing CST axons. The recordings were sampled at 30KS/s and recorded using an integrated data acquisition system (Smartbox; NeuroNexus Technologies). Paired t-test was conducted to determine baseline change in firing rate before and after the light stimulation. Units that exhibited mean firing rate change greater than 2 spikes/sec and a significant difference (p<.0001) were classified as exhibiting laser-evoked activity in this analysis.

### Axon growth quantification

Spinal cord sections were cut using a vibratome (Leica VT100S) at 50 µm thickness. Axon growth was quantified by a blinded observed from sagittal sections imaged on Nikon A1 Confocal Microscope and analyzed with Advanced Research software. Fluorescent axons that intersected virtual lines at set distances from the injury site were quantified and normalized to total fluorescent axons counted in the medullary pyramids

