## Supplemental Methods and Figures for "Restoration of Direct Corticospinal Communication Across Sites of Spinal Injury"

### Extended Methods

*AAV preparation*

Mouse *KLF6* (accession BC020042), was purchased from Dharmacon and the open reading frame along with 5' and 3' UTRs cloned into an AAV-compatible plasmid with CMV reporter (pAAV-MCS, Stratagene) using standard PCR amplification as in (1, 2). Production of AAV9-KLF6, AAV9-EBFP and AAV9-EGFP (1), and AAV9–CaMKII–ChR2(H134R)–EYFP (3–5) was performed at the University of North Carolina Viral Vector Core.

*Cortical AAV injections and spinal injuries*

All experiments were conducted following protocols approved by the Marquette University Animal Care and Use Committee, in accordance with the National Institutes of Health Guide for the Care and Use of Laboratory Animals. Female C57/Bl6 mice were housed in a temperature and humidity controlled vivarium with food and water available *ad libitum*. For cortical delivery of virus, animals were anesthetized with ketamine (100mg/kg) and xylazine (10mg/kg) and placed in a stereotaxic frame (Stoelting Mouse Adaptor), craniotomy performed, and 0.3µL viral particles ($1 \times 10^{13}$/ml) were delivered by Hamilton syringe and Stoelting QSI infusion pump, (0.03 µL/min) to 3 sites located 1.2-1.5 mm lateral (left) and 0.2-0.7 anterior from Bregma and 0.55 mm below the surface of the brain. The needle was then withdrawn, and the skin closed with staples. For spinal cord injury animals were anesthetized using a mixture of ketamine/xylazine, mounted into a custom spine stabilizer, and the dorsal surface of the spinal cord exposed between C4/C5. Using stereotaxic control, a tungsten wire knife (McHughMilieux, M120) was situated 950µm to the right of the midline, the wire extended to the midline, and lowered until the blade reached a depth of 1.1mm. The knife was then raised to the surface of the spinal cord while maintaining downward pressure on the dorsal surface to ensure a complete lesion. This resulted in an injury of at least 1.1mm depth, extending 1mm lateral from the midline.

*Stem cell preparation and transplantation*

Timed pregnancies between B6.129(Cg)-Gt(ROSA)26Sortm4(ACTB-tdTomato,-EGFP)Luo/J transgenic male mice and wildtype females produced E12 embryos that coincided with the one week post-injury time point. Pregnant mothers were anesthetized by exposure to isoflurane, decapitated, and embryos were removed to ice-cold PBS. Cervical and upper thoracic spinal cord were dissected, washed 3X in ice-cold Ca+/Mg+-free HBSS (Gibco 14175), dissociated by ~10X passage through a fire-polished pipette, brought to 10ml in Neurobasal media, and pelleted (650G, 2min). The pellet was resuspended in 1ml Neurobasal media, passed through a 40µm filter (Falcon 352340), and brought to 1 million cells/ml in ice-cold Neurobasal media. For each animal, 500,000 cells in two tubes were pelleted at 650G, 2min, and resuspended in 1.5µl of thrombin (50U/ml, Sigma) or fibrinogen (50mg/ml, Sigma) with BDNF (Peprotech, 50ng/ul), VEFG (Peprotech, 10ng/ul), bFGF(Peprotech, 10ng/ul) and MDL28170VEGF (Sigma, 50µm).

To deliver stem cells to the spinal cord, mice were anesthetized with ketamine/xylazine and mounted in a specialized frame to immobilize the spinal cord and re-expose the site of spinal injury. Using a Picospritzer III (Parker Hannifin Corp) interfaced with a pulled glass pipette, cells mixed in fibrinogen were injected to five locations within and adjacent to the injury, with three injections distributed across the medial/lateral extent of the injury, one site 500µm rostral and another 500µm caudal. Approximately 100,000 cells were injected in 100nl to each location, using a rapid series of picospritzer pulses that expelled about 10nl per pulse. Immediately afterward, the thrombin/cell mixture was injected to identical locations, with similar cell concentrations and rates of expulsion. After the transplantation, the muscles were sutured back in place, and the skin was closed using surgical staples.

*Behavioral testing*

In the horizontal ladder task, mice traversed a ladder 30cm in length, with rungs separated by 1cm, and suspended between 2 plates of plexiglass at a height of 10cm. Prior to injury, animals were pre-trained and then tested in 4-crossing sessions, with percent error quantified on the last three runs. Animals were similarly tested 4 and 8 weeks post-injury. At 8 weeks, animals were also tested on ladders with unevenly spaced rungs, created by removing a randomized subset of rungs from the original ladder. All trials were video-recorded and errors were analyzed by blinded observers. Errors were defined as slips by the right forelimb (wrist breaks plane of ladder) and quantified as percentage of total steps by the right forelimb. For pellet retrieval testing, animals were food restricted to maintain 80% of their body weight and placed in staircase pellet retrieval apparati (Lafayette Instruments, Lafayette IN) with two high fat diet 20mg pellets (BioServ, Flemington NJ) placed on each of eight steps of increasing depth. Animals were allowed 30 minutes to retrieve as many pellets as possible. Animals were pre-trained by daily exposure to the task for at least two weeks. After injury, animals were tested twice weekly, and the total number of pellets that were retrieved and displaced during both sessions was recorded (6).

*Optical stimulation/ Single-unit extracellular recordings*

Optogenetic stimulation and single-unit extracellular recordings were performed as previously described. Briefly, mice were completely anesthetized using urethane (IP, 10mg/kg) and their spinal cords between C4 and C5 were exposed as for spinal surgery, above. The center-point of the spinal injury was visually identified. Using stereotactic guidance (David KOPF) a 32-channel silicon electrode (NeuroNexus Technologies) was inserted 1000µm deep into a series of recording locations (at the injury/stem cell center-point, and 500µm and 1000µm rostral and caudal to this location). At all locations a 473 nm laser was focused via collimator on the point of electrode insertion to stimulate terminal firing of ChR2-expressing CST axons. Output power (5 - 20mW) from the tip of the fiber was measured using an optical power meter (Thorlabs, Inc, PM130D) and was adjusted to maximize the effectiveness of stimulation. Pulses were generated using TTL modulation (Arbitrary Waveform Generator, 20 MHz, Agilent 33220A). Stimulation consisted of six pulse trains, 15 seconds apart, with each train comprised of 3 pulses, 1 second each with 1 second intervals.

*Data acquisition and analysis*

Electrode signals were relayed via a smart-link head stage and then amplified, digitized, and stored in an integrated unit (Smartbox; NeuroNexus Technologies). Signals were sampled at 30 KS/s, with digital to analog converter (DAC) high pass filter settings were set to 250 Hz and the Notch filter to 60 Hz, and exported to Spike2 v8.00 (CED) (5). Discrimination of individual waveforms corresponding to the activity of an individual neuron was accomplished with principal component analysis using Offline Sorter (Plexon Inc). Only biphasic waveforms indicative of somatic recordings were observed. To quantify and statistically compare firing rates, peri-event raster histograms (100ms bins) surrounding each stimulation event were generated using Neuroexplorer 4.126 (Nex Technologies). Each histogram was divided into 18 baseline epochs (1 sec prior to each laser onset) and 18 matching stimulation epochs timelocked to laser duration (1 sec). For each unit, a paired t-test ($\alpha = .05$) was conducted comparing baseline and stimulation firing rate and units that exhibited mean firing rate change greater than 2 spikes/sec and a significant difference ($p < .0001$) in this analysis were classified as exhibiting laser-evoked activity. $\chi^2$ tests were used to compare the proportion of light-responsive cells across treatments.

*Histology and quantification of axon growth*

Immediately after electrophysiological recordings, animals were euthanized with CO2 and perfused with 4% paraformaldehyde. Cortices, medullas, and spinal cords were removed, fixed in 4% PFA overnight at 4°C, and embedded in 6% gelatin (Sigma) for sectioning. Cortex and medulla were sectioned in the coronal plane, and cervical spinal cord spanning 2mm rostral to 4mm caudal to the injury was sectioned in the horizontal plane. All sections were cut on a Leica VT100S Vibratome at 50µm, free-floating in PBS with 0.02% $NaN_3$. Three spinal cord sections per animal were imaged using a Nikon A1 confocal microscope. Spinal cord images were taken at 20x magnification and maximum intensity projections were created from 6 Z-planes that spanned 25µm of tissue depth. Virtual lines were created at 200µm intervals from the lesion site, oriental orthogonally to the rostral/caudal axis of the spinal cord. The number of EYFP+ axons that intersected each line was quantified by a blinded observer. Similar quantification was performed using measurement lines positioned relative to the point of CST retraction from the injury site, or from the caudal-most edge of the stem cell grafts. In the latter case, measurement lines followed the shape of the graft/host interface and were spaced at 100µm intervals. Axon counts were normalized to the total number of EYFP+ axons detected in the medullary pyramids. Medullary counts were obtained using a Nikon Eclipse Ti-S inverted microscope at 60x magnification, subsampling three regions (4000µm$^2$ each) and then extrapolating based on total cross-sectional area of the pyramid. All spinal and medulla counts were made by blinded observers. Exclusion criteria for horizontal spinal cord sections included low EYFP signal intensity in CST axons that was too dim to reliably distinguish from background fluorescence, and incomplete injuries as determined by the presence of linear EYFP+ axons that extended through tdTomato- tissue (typically spared dorsolateral CST axons. Full animal mortality and exclusions are provided in **Supplemental Table 1.**

| Table 1 : Animal numbers and exclusions | | | | | | | |
|---|---|---|---|---|---|---|---|
| Group | Starting animals | Mortality | Loss in tissue processing | Failed Stem cell Integration or overgrowth | Incomplete Injury | Failed CST labelling | *Final N* |
| **No Stem Cells, Control virus** | 10 | 2 | 2 | NA | 0 | 1 | **5** |
| **No Stem Cells, KLF6** | 10 | 2 | 2 | NA | 0 | 0 | **6** |
| **Stem Cells, Control virus** | 18 | 3 | 1 | 0 | 2 | 0 | **12** |
| **Stem Cells, KLF6** | 19 | 4 | 0 | 1 | 2 | 0 | **12** |
| **Stem Cells, Retro KLF6** | 10 | 2 | 0 | 0 | 0 | 0 | **8** |

Supplemental Table 1: Starting animal numbers, mortality, and exclusions for each experimental group.

Supplemental Figure 1

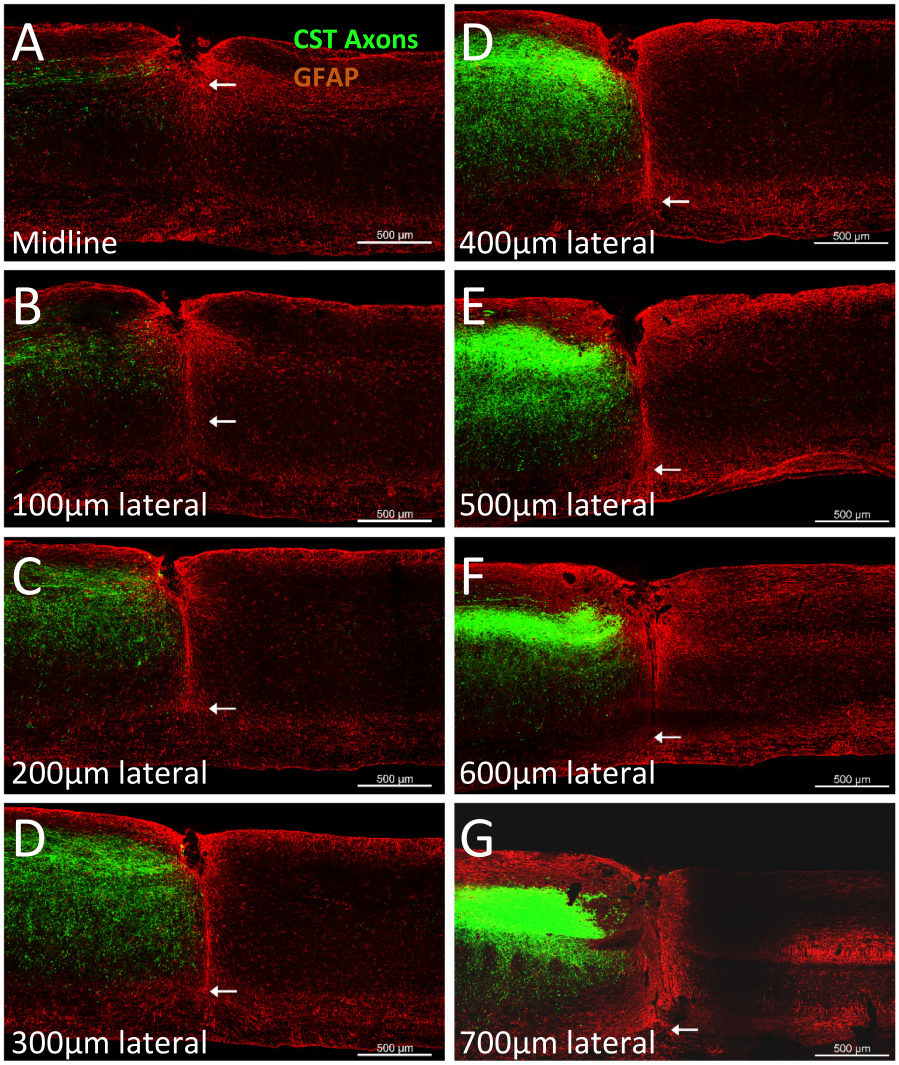


**Supplemental Figure 1**. Confirmation of injury depth. Adult mice received a wireknife injury between C4 and C5 spinal cord, and cortical injection of AAV-EGFP to label corticospinal tract neurons. Eight weeks later, sagittal sections of cervical spinal cord were stained for GFAP (red) and imaged to determine the depth of transection. A-H show an example of a sagittal series of the right side of the spinal cord, with arrows indicating the depth of the injury. This analysis confirms complete severing of the dorsal CST (A) and dorsal-lateral CST tracts (H), and that the injury extends 1.2mm from the dorsal surface of the spinal cord and extends throughout the grey matter.

Supplemental Figure 2

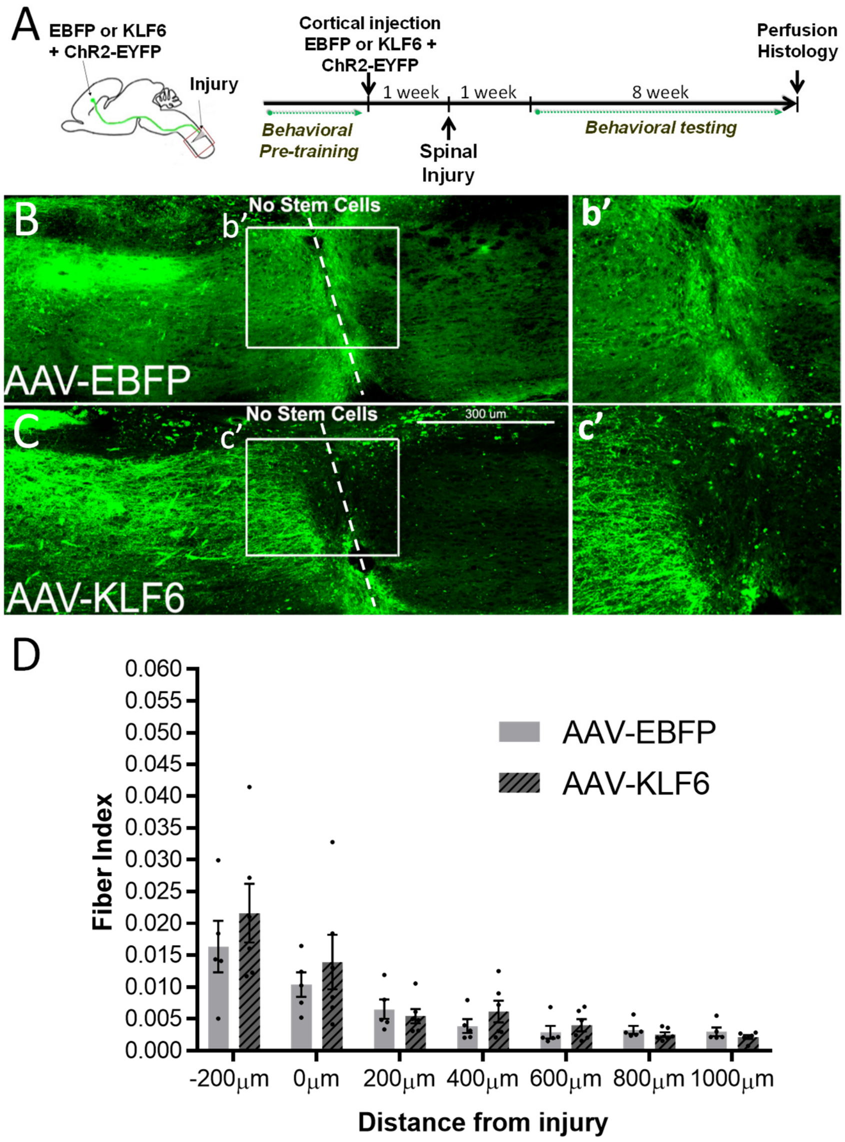


**Supplemental Figure 2.** After deep cervical injury, KLF6 treatment does not enhance CST regeneration beyond the injury site. A) Indicates the sites and timing of spinal transections and cortical injection of AAV-EBFP or AAV-KLF6 in adult mice. (B,C) Horizontal sections of cervical spinal cord, with CST axons labeled green. In both EBFP control animals and KLF6-treated animals, CST axons extend minimally into or beyond the injury (dotted line). D) Quantification of EGFP+ CST profiles shows no significant difference between EBFP and KLF6-treated animals in the number of axons that approach or grow beyond the spinal injury site. Each dot represents average axon counts from an individual animal, bars indicate average of the treatment group. N=5 control, 6 KLF6. ($p>.05$, repeated measures ANOVA).

Supplemental Figure 3

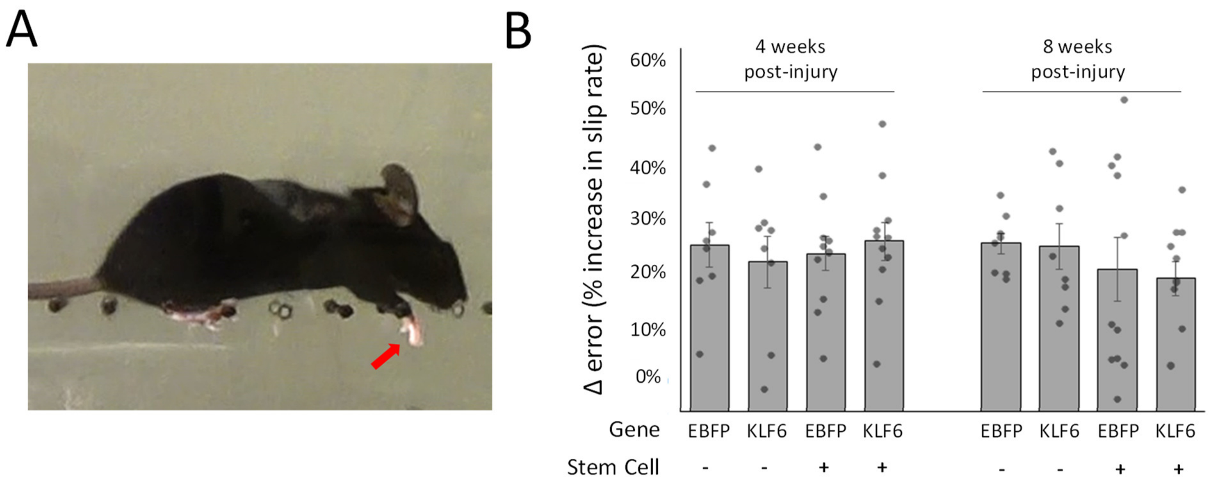


Supplemental Figure 3. C57/Bl6 mice received cervical dorsal hemisection and cortical injection of AAV-EBFP or AAV-KLF6, followed one week later by stem cell grafts. At 4 or 8 weeks post-injury animals were tested on the horizontal ladder task, quantifying the number of errors in the right forelimb. (A) Forelimb steps in which the wrist descended below the plane of the ladder were counted as errors. (B) The percent of errors for each animal, normalized to the average number of errors made by the same animal prior to injury, was determined. Stem cell treatment, KLF6 treatment, and the two combined had no effect on the error rate ($p>.05$, 2-way ANOVA, $N\geq7$ animals per group).
